# Genetic dissection of an ancient divergence in yeast thermotolerance

**DOI:** 10.1101/220913

**Authors:** Carly V. Weiss, Jeremy I. Roop, Rylee K. Hackley, Julie Chuong, Igor V. Grigoriev, Adam P. Arkin, Jeffrey M. Skerker, Rachel B. Brem

**Affiliations:** Department of Plant and Microbial Biology, UC Berkeley, Berkeley, CA.; Current address: Fred Hutchinson Cancer Research Center, Seattle, WA.; Buck Institute for Research on Aging, Novato, CA.; Current address: Duke University Program in Genetics & Genomics, Duke University, Durham, NC.; US Department of Energy Joint Genome Institute, Walnut Creek, CA.; Department of Bioengineering, UC Berkeley, Berkeley, CA and Lawrence Berkeley National Laboratory, Berkeley, CA.

## Abstract

Some of the most unique and compelling survival strategies in the natural world are fixed in isolated species. To date, molecular insight into these ancient adaptations has been limited, as classic experimental genetics has focused on interfertile individuals in populations. Here we use a new mapping approach, which screens mutants in a sterile interspecific hybrid, to identify eight housekeeping genes that underlie the growth advantage of *Saccharomyces cerevisiae* over its distant relative *S. paradoxus* at high temperature. Pro-thermotolerance alleles at these mapped loci were required for the adaptive trait in *S. cerevisiae* and sufficient for its partial reconstruction in *S. paradoxus*. The emerging picture is one in which *S. cerevisiae* improved the heat resistance of multiple components of the fundamental growth machinery in response to selective pressure. This study lays the groundwork for the mapping of genotype to phenotype in clades of sister species across Eukarya.

---

Geneticists since Mendel have sought to understand how and why wild individuals differ. Studies toward this end routinely test for a relationship between genotype and phenotype via linkage or association^1^. These familiar approaches, though powerful in many contexts, have an important drawback—they can only be applied to interfertile members of the same species. This rules out any case in which an innovation in form or function evolved long ago and is now fixed in a reproductively isolated population.

As organisms undergo selection over long timescales, their traits may be refined by processes quite different from those that happen early in adaptation^2,3^. We know little about these mechanisms in the wild, expressly because when the resulting lineages become reproductively incompatible, classic statistical-genetic methods cannot be used to analyze them^4^. To date, the field has advanced largely on the strength of candidate-based studies that implicate a single variant gene in an interspecific trait^5,6^, with the complete genetic architecture often remaining unknown. Against the backdrop of a few specialized introgression^7–10^ and molecular-evolution^11^ techniques available in the field, dissection of complex trait differences between species has remained a key challenge.

Here we develop a new genetic mapping strategy, based on the reciprocal hemizygosity test^12,13^, and use it to identify the determinants of a difference in high-temperature growth between isolated *Saccharomyces* yeast species. We validate the contributions of the mapped loci to the thermotolerance trait, and we investigate their evolutionary history.

## Results

### Species differences in thermotolerance

At high temperature, the yeast *Saccharomyces cerevisiae* grows qualitatively better than other members of its clade^14–16^, including its closest relative, *S. paradoxus*, from which it diverged ~5 million years ago^17^. In culture at 39°C, *S. cerevisiae* doubled faster than *S. paradoxus* and accumulated more biomass over a timecourse, a compound trait that we call thermotolerance. The magnitude of differences in thermotolerance between species far exceeded that of strain variation within each species (Figure 1), whereas no such effect was detectable at 28°C (Figure S1). The failure by *S. paradoxus* to grow to high density at 39°C was, at least in part, a product of reduced survival relative to that of *S. cerevisiae*, as cells of the former were largely unable to form colonies after heat treatment (Figure S2). In microscopy experiments, *S. paradoxus* cells were almost uniformly visible as large-budded dyads after 24 hours at 39°C (Figure S3), suggestive of defects late in the cell cycle as a proximal cause of death^18^; no such tendency was apparent in *S. cerevisiae* at high temperature, or in either species at 28°C (Figure S3).

**Figure 1.**
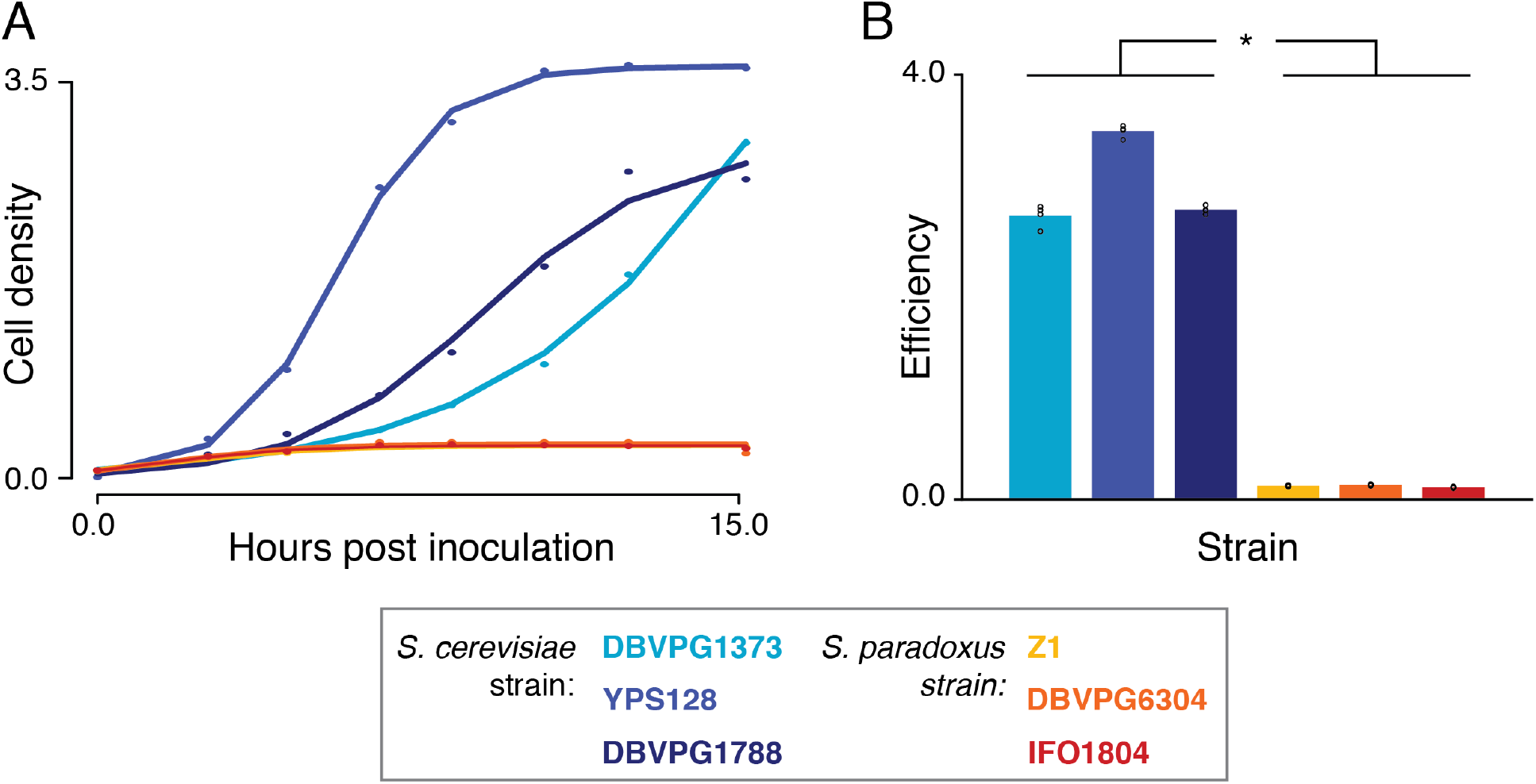
*S. cerevisiae* grows better at high temperature than *S. paradoxus*. **A**, Each point reports cell density (OD_600_) of the indicated wild isolate of *S. cerevisiae* (blue) or *S. paradoxus* (orange) over a timecourse of growth at 39°C. Each solid line shows a logistic population growth model fit to the respective cell density measurements. **B**, Each bar reports mean efficiency (*n* = 4) of the indicated strain at 39°C, defined as the difference between cell density at 24 hours of growth and that at the time of inoculation. *, *p* = 3.78×10^−11^; individual measurements are reported as open circles.

### Massively parallel reciprocal hemizygosity testing by RH-seq

We set out to dissect the genetic basis of *S. cerevisiae* thermotolerance, using a genomic implementation of the reciprocal hemizygote test^12,13^ (Figure 2A). For this purpose, we first mated *S. cerevisiae* strain DBVPG1373, a soil isolate from the Netherlands, with *S. paradoxus* strain Z1, an English oak tree isolate. The resulting sterile hybrid had a thermotolerance phenotype between those of its purebred parents (Figure S4). In this hybrid background we generated hemizygote mutants using a plasmid-borne, selectable PiggyBac transposon system^19^. We cultured the pool of mutants in bulk for ~7 generations at 39°C and, separately, at 28°C. From cells in each culture we sequenced transposon insertion locations^20^ as a readout of the genotypes and abundance of mutant hemizygote clones present in the selected sample. In these sequencing data, at each of 4888 genes we detected transposon mutant clones in both species’ alleles in the hybrid (Figure S5), with transposon insertions distributed in a largely unbiased manner across the genome (Figure S6). For a given gene, we tabulated the abundances of mutants whose transposon insertion fell in the *S. cerevisiae* allele of the hybrid, after high-temperature selection relative to the 28°C control, and we compared them to the abundance distribution of mutants in the *S. paradoxus* allele (Figure 2A). Any difference in abundance between these reciprocal hemizygote cohorts can be ascribed to variants between the retained alleles at the respective locus; we refer to the comparison as reciprocal hemizygosity analysis via sequencing (RH-seq). Integrating this approach with a quality-control pipeline (Figure S5), in a survey of 3416 high-coverage genes we identified 8 top-scoring hits (false discovery rate 0.01; Figure 2B). At each such locus, disruption of the *S. cerevisiae* allele in the hybrid was associated with low clone abundance after selection at 39°C relative to 28°C (Figure 2B), reflecting a requirement for the *S. cerevisiae* allele for thermotolerance. None of the genes mapped by RH-seq had a known role in the yeast heat shock or stress responses. All were annotated as housekeeping factors: *ESP1, DYN1, MYO1, CEP3, APC1*, and *SCC2* function in chromosome segregation and cytokinesis, and *AFG2* and *TAF2* in transcription/translation.

**Figure 2.**
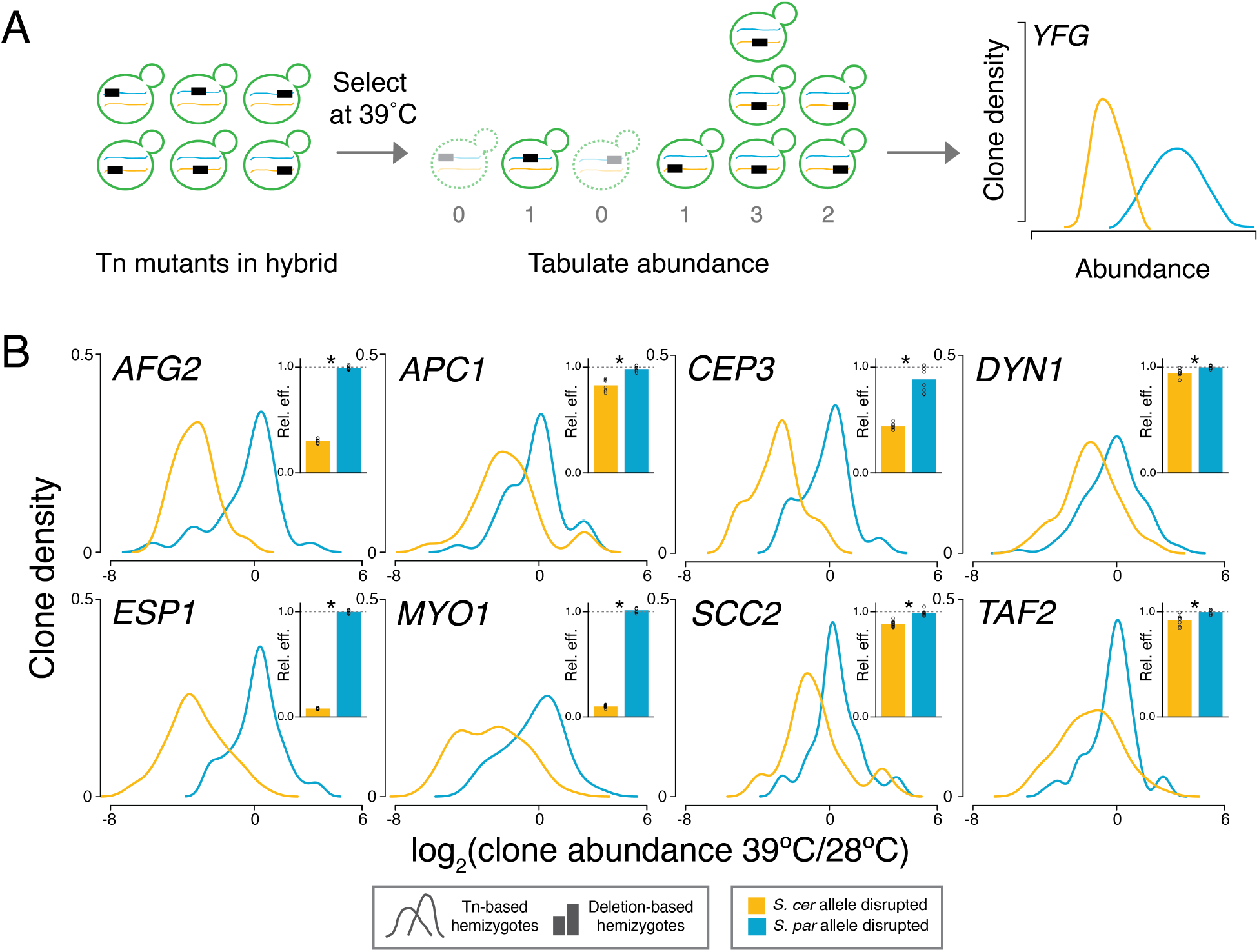
Mapping thermotolerance by RH-seq. **A**, A transposon (rectangle) disrupts the allele from *S. cerevisiae* (blue) or *S. paradoxus* (orange) of a gene (*YFG*) in an interspecific hybrid (green). Clones lacking the pro-thermotolerance allele grow poorly at 39°C (dashed outlines), as measured by sequencing and reported in smoothed histograms (traces, colored to indicate the species’ allele that is not disrupted). **B**, Each panel reports results from one RH-seq hit locus. In the main figure of a given panel, the x-axis reports the log_2_ of abundance, measured by RH-seq after selection at 39°C, of a clone harboring a transposon insertion in the indicated species’ allele, relative to the analogous quantity for that clone from selection at 28°C. The *y*-axis reports the proportion of all clones bearing insertions in the indicated allele that exhibited the abundance ratio on the *x*, as a kernel density estimate. In insets, each bar reports the relative efficiency, calculated as the mean growth efficiency at 39°C (*n* = 8-12) of the indicated targeted-deletion hemizygote measured in liquid culture assays, normalized to the analogous quantity for the wild-type hybrid. *, *p* ≤ 0.002; individual measurements are reported as open circles.

To evaluate the role in thermotolerance of genes that emerged from RH-seq, we first sought to verify that growth differences between hemizygotes at a given locus were the consequence of allelic variation and not an artifact of our genomic approach. As a complement to our analysis of transposon mutants, for each RH-seq hit gene we engineered hemizygotes by targeted deletion of each species’ allele in turn in the hybrid. In growth assays, the strain lacking the *S. cerevisiae* allele at each gene grew poorly at high temperature (Figure 2B), with little impact at 28°C (Figure S7), consistent with fitness inferences from RH-seq. And in each case, the *S. paradoxus* allele made no contribution to the phenotype of the hybrid, since deleting it had no effect (Figures 2B and S7). We conclude that the loci emerging from RH-seq represent bona fide determinants of thermotolerance in the hybrid.

### Validation of RH-seq gene hits in purebreds

We expected that variation at our RH-seq hits, though mapped by virtue of their impact in the hybrid, could also explain thermotolerance differences between purebred species. As a test of this notion, for each mapped gene in turn, we replaced the two copies of the endogenous allele in each purebred diploid with the allele from the other species. Growth assays of these transgenics established the *S. cerevisiae* allele of each locus as necessary or sufficient for biomass accumulation at 39°C, or both: thermotolerance in the *S. cerevisiae* background was compromised by *S. paradoxus* alleles at 7 of the 8 genes and, in *S. paradoxus*, improved by *S. cerevisiae* alleles at 6 of 8 loci (Figure 3). Allele replacements had little impact on growth at 28°C (Figure S8). These trends mirrored the direction of locus effects from hemizygotes in the hybrid, though the magnitudes were often different. Most salient were the small effect sizes in *S. paradoxus* relative to other backgrounds, indicative of strong epistasis in this poorly-performing species (Figure S9). Thus, the loci mapped by RH-seq in an interspecies hybrid contribute causally to thermotolerance in purebreds, with effect sizes that depend on the context in which they are interrogated.

**Figure 3.**
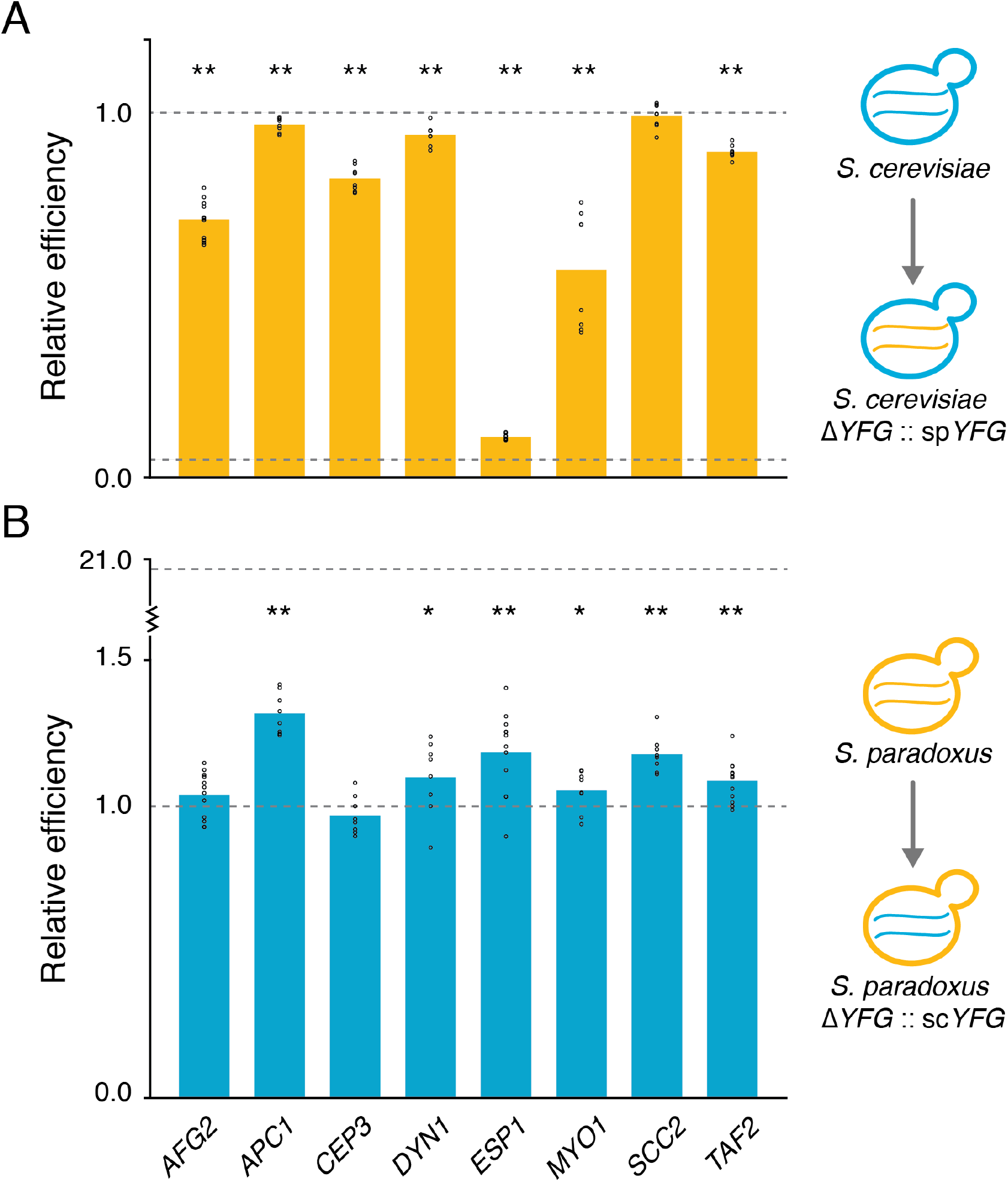
*S. cerevisiae* thermotolerance alleles are necessary and sufficient for growth at high temperature. **A**, Each bar reports mean growth efficiency at 39°C, measured in liquid culture assays (n = 6-12), of an *S. cerevisiae* strain harboring the *S. paradoxus* allele at the indicated RH-seq hit locus, relative to the analogous quantity for wild-type *S. cerevisiae*. **B**, Data are as in **A**, except that each bar reports results from a *S. paradoxus* strain harboring the *S. cerevisiae* allele at the indicated locus, normalized to wild-type *S. paradoxus*. In a given panel, the top and bottom dotted lines report the relative efficiency of wild-type *S. cerevisiae* and *S. paradoxus*, respectively. *, *p* ≤ 0.036; **, *p* ≤ 0.001. Individual measurements are reported as open circles.

Avid growth at high temperature is a defining characteristic of *S. cerevisiae* as a species, relative to other Saccharomycetes (refs. 14-16 and Figure 1). In principle, the loci mapped by RH-seq could be unique to the genetic architecture of thermotolerance in our focal *S. cerevisiae* strain, DBVPG1373, or be part of a mechanism common to many *S. cerevisiae* isolates. In support of the latter model, transgenesis experiments showed that a diverse panel of *S. cerevisiae* isolates all harbored alleles conferring modest but significant growth benefits at high temperature, and alleles from multiple *S. paradoxus* isolates were deleterious (Figure S10A-B). We detected no such impact at 28°C (Figure S10A-B). Similarly, we found elevated sequence divergence from *S. paradoxus* to be a shared feature of *S. cerevisiae* strains at the loci mapped by RH-seq (using the absolute divergence measure D_xy_; Figure S10C). These findings indicate that the *S. cerevisiae* population accumulated divergent, pro-thermotolerance alleles at the loci mapped by RH-seq, consistent with a role in the trait for these genes across the species. Likewise, in the yeast phylogeny, RH-seq hit genes were distinguished by accelerated evolution along the branch leading to *S. cerevisiae*, as expected if the ancestral program has been conserved among the other species in the clade (Figure S10C).

## Discussion

In this work, we have developed the RH-seq method for genome-wide mapping of natural trait variation, and we have used it to elucidate the genetics of thermotolerance in reproductively isolated yeasts. Growth at high temperature is likely a derived character in S. cerevisiae^14–16^, and the mechanism by which evolution built the trait, after the split from *S. paradoxus*, has remained unknown. In pursuing the genetics of this putative ancient adaptation, we complement studies of younger, intra-specific variants that erode thermotolerance, in the few *S. cerevisiae* strains that have lost the trait relatively recently^12,21^. We have sought to shed light on more ancient evolutionary events by considering *S. paradoxus* as a representative of the ancestral state, to which thermotolerant *S. cerevisiae* can be compared.

Using this approach, we have mapped eight loci at which *S. cerevisiae* alleles are necessary and sufficient for thermotolerance. Six of these genes are essential for growth in standard conditions^22^, and all eight contribute to fundamental growth processes. *ESP1, DYN1, CEP3, APC1, MYO1*, and *SCC2* mediate mitotic spindle assembly, chromatid cohesion and separation, cytokinesis, and mitotic exit; *AFG2* regulates the release of maturation factors from the ribosome; and *TAF2* encodes a TFIID subunit. In each case, our growth experiments in the interspecific hybrid have shown that the *S. paradoxus* allele acts as a hypomorph at high temperature. Our work leaves unanswered exactly how heat-treated *S. paradoxus* cells die in the absence of these functions, though their large-budded morphology strongly suggests regulated arrest or stochastic failure late in the cell cycle. Given that some but not all RH-seq hit loci have roles in mitosis, it is likely only one of the choke points at which *S. paradoxus* alleles are a liability at high temperature. Assuming that these heat-sensitive alleles also littered the genome of the common ancestor with *S. cerevisiae*, thermotolerance would have evolved along the *S. cerevisiae* lineage by resolving each of them, boosting the heat resistance of many housekeeping processes. Such a mechanism would dovetail with the recent finding that, across species, the limiting temperature for cell growth correlates with the melting temperatures of a subset of essential proteins^23^.

These insights into the evolution of a complex yeast trait serve as a proof of concept for RH-seq. To date, the reciprocal hemizygosity test has led to landmark discoveries in a candidate-gene framework, confirming the effects of variation at a given locus identified by other means^12,13^. Schemes to scale up the test have generated a genome’s worth of hemizygotes from deletion-strain purebreds, which tend to harbor secondary mutations that come through screens as false positives^24,25^. As such, a key advantage of RH-seq is that we carry out mutagenesis in the hybrid, which ensures coverage of essential genes and obviates the use of mutation-prone null genotypes. Furthermore, any secondary mutations that do arise in a given hemizygote clone should have little influence in RH-seq mapping, because deep mutagenesis generates many independent clones per gene that are analyzed together. One important caveat of RH-seq, as in single-gene reciprocal hemizygote tests, is the assumption that no epistasis unique to the hybrid will mask the effects of loci underlying a trait difference of interest between the parents. In our case study, the genetic architecture of thermotolerance in the hybrid did bear out as relevant for the purebreds, albeit with locus effect sizes that varied across the backgrounds. More dramatic discrepancies may be particularly likely when the hybrid has a heterotic (*i.e*. extreme) phenotype and is a poor model for the genetics of the parents^26^. The choice of a non-heterotic hybrid in which to pursue RH-seq would be analogous to classical linkage mapping in a cross whose progeny have, on average, phenotypes that are intermediate between those of the parents.

In fact, although we have focused here on ancient divergence, the RH-seq method would be just as applicable to individuals within a species, as a high-resolution alternative to linkage analysis. We thus anticipate that RH-seq will accelerate the mapping of genotype to phenotype in many systems, whether the parents of a cross are closely related or members of a species complex that have been locally adapting for millions of years.

## Materials and Methods

### Strains

Strains used in this study are listed in Table S1. Homozygous diploid strains of *S. cerevisiae* and *S. paradoxus* used as parents of the interspecific hybrid, and as the backgrounds for allele-swap experiments, were homothallic DBVPG1373 and Z1, respectively. In the case of the hybrid parents, each strain was rendered homozgyous null for *URA3* via homologous recombination with a HYGMX cassette, then sporulated; a given mated spore from a dissected tetrad was grown up into a diploid that was homozygous null at *URA3* and tested for the presence of both genomes by PCR with species-specific primers.

### piggyBac transposon machinery

For untargeted, genome-scale construction of reciprocal hemizygotes in the *S. cerevisiae* x *S. paradoxus* hybrid, we adapted methods for piggyBac transposon mutagenesis^19^ to develop a system in which the transposon machinery was borne on a selectable and counter-selectable plasmid lacking a centromere. We constructed this plasmid (final identifier pJR487) in three steps. In step I we cloned the piggyBac transposase enzyme gene driven by the *S. cerevisiae TDH3* promoter (from plasmid p3E1.2, a gift from Malcolm Fraser, Notre Dame) into plasmid pJED104 (which contains *URA3*, an ARS, and the *CEN6* locus, and was a gift from John Dueber, UC Berkeley). For this cloning, the amplification used a forward and reverse primer containing a BamHI and XhoI site, respectively, that upon restriction digest yielded sticky ends for ligation to recipient BamHI and XhoI sites in digested pJED104. We used the resulting plasmid as input into step II, removal of the *CEN6* sequence: we first amplified the entire plasmid with primers that initiated outside of *CEN6* and were directed away from it, and contained reciprocally complementary NheI sites; sticky ends of the linear PCR product were then ligated together for re-circularization. We used the resulting plasmid as input into step III, the cloning in of a construct comprised of the KANMX cassette flanked by long terminal arms (328bp and 361bp) from the piggyBac transposon. We first amplified KANMX from pUG6^27^ and each transposon arm from p3E1.2, using primers that contained overlapping sequence on the fragment ends that would ultimately be the interior of the construct, and XbaI sites on the fragment ends that would ultimately be the 5’ and 3’-most ends of the construct. We stitched the three fragments together by overlap extension PCR, digested the resulting construct and the plasmid from step II with XbaI, and annealed sticky ends of the two to yield the final pJR487 plasmid.

### Untargeted hemizygote construction via transposon mutagenesis

For mutagenesis, pJR487 was gigaprepped using a column kit (Zymo Research) to generate ~11 mg plasmid. To prepare for transformation, JR507 (the *S. cerevisiae* DBVPG1373 x *S. paradoxus* Z1 hybrid) was streaked from a −80°C freezer stock onto a yeast peptone dextrose (YPD, 1% yeast extract [BD], 2% yeast peptone [BD], 2% D-glucose [Sigma]) agar plate and incubated for 2 days at 26°C. A single colony was inoculated into 100 mL YPD and shaken at 28°C, 200rpm for ~24 hours. The next day, we transferred cells from this pre-culture, and YPD, to each of four 1 L flasks at the volumes required to attain an optical density at 600 nm (OD_600_) of 0.2 in 500 mL each. We cultured each for 6 hours at 28°C with shaking at 200rpm. Two of these cultures were combined into 1 L of culture and two into a separate 1 L, and each such culture was subjected to transformation (for a total of two transformations) as follows. The 1 L was split into twenty 50-mL conical tubes. Each aliquot was centrifuged and washed with water and then with 0.1 M lithium acetate (LiOAc, Sigma) mixed with 1X Tris-EDTA buffer (10 mM Tris-HCl and 1.0 mM EDTA); after spin-down, to each tube was added a solution of 0.269 mg of pJR487 mixed 5:1 by volume with salmon sperm DNA (Invitrogen), and then to each was added 3 mL of 39.52% polyethylene glycol, 0.12M LiOAc and 1.2X Tris-EDTA buffer (12 mM Tris-HCl and 1.2 mM EDTA). Tubes were rested for 10 minutes at room temperature, then heat-shocked in a water bath at 39°C for 26 minutes. Cells from all 20 tubes were then combined. We transferred cells from this post-transformation culture, and YPD, to each of three 1 L flasks at the volumes required to attain an OD_600_ of ~0.35-4 in 500 mL. Each such culture was recovered by shaking at 28°C and 200 rpm for 2 hours. G418 (Geneticin, Gibco) was added to each at a concentration of 300 μg/mL to select for those cells which had taken up the plasmid, and cultures were incubated with 200 rpm shaking at 28°C for two days until each reached an OD_600_ of ~2.3. All six such selected cultures across the two transformations were combined. We transferred cells from this combined culture, and YPD + G418 (300 ug/mL), to each of two 1 L flasks at the volumes required to attain an OD_600_ of 0.2 in 500 mL each. We cultured each flask at 28°C and 200 rpm shaking overnight until reaching an OD_600_ of 2.18 and combined the two cultures again to yield one culture. To cure transformants of the pJR487 URA+ plasmid, we spun down a volume of this master culture and resuspended in water with the volume required to attain a cell density of 1.85 OD_600_ units/mL. 12 mL of this resuspension were plated (1 mL per 24.1cm x 24.1cm plate) onto plates containing complete synthetic media with 5-fluoroorotic acid (5-FOA) [0.2% drop-out amino acid mix without uracil or yeast nitrogen base (YNB) (US Biological), 0.005% uracil (Sigma), 2% D-glucose (Sigma), 0.67% YNB without amino acids (Difco), 0.075% 5-FOA (Zymo Research)]. After incubation at 28°C to enable colony growth, colonies were scraped off all 12 plates and combined into water at the volume required to attain 40 OD_600_ units per 900 μL, yielding the final transposon mutant hemizygote pool. This was aliquoted into 1 mL volumes with 10% DMSO and frozen at −80°C.

### Thermotolerance phenotyping via selection of the hemizygote pool

One aliquot of the pool of transposon mutant hemizygotes in the JR507 *S. cerevisiae* DBVPG1373 x *S. paradoxus* Z1 hybrid background was thawed and inoculated into 150 mL of YPD in a 250 mL flask, and cultured for 7.25 hours at 28°C, with shaking at 200 rpm. We used this timepoint as time zero of our thermotolerance experiment, and took four aliquots of 6.43 mL (7 OD units) as technical replicates for sequencing of transposon insertion positions (see below). 9.19 mL of the remaining culture was back-diluted to an OD_600_ of 0.02 in a total of 500 mL YPD in each of six 2L glass flasks for cultures that we call selections; three were grown at 28°C and three at 39°C (shaking at 200 rpm) until an OD_600_ of 1.9-2.12 was reached, corresponding to about 6.5 doublings in each case. Four cell pellets of 7 OD_600_ units each were harvested from each of these biological replicate flasks, for sequencing as technical replicates (see below). In total, 28 pellets were subjected to sequencing: 4 technical replicates from the time-zero culture; 3 biological replicates, 4 technical replicates each, from the 28°C selection; and 3 biological replicates, 4 technical replicates each, from the 39°C selection.

### Tn-seq library construction

To determine the abundance of transposon mutant hemizygote clones after selection, we first sequenced transposon (Tn) insertions as follows. Each cell pellet from a time zero or selection sample (see above) was thawed on ice, and its genomic DNA (gDNA) was harvested with the ZR Fungal/Bacterial DNA MiniPrep kit (Zymo Research). gDNA was resuspended in DNA elution buffer (Zymo) pre-warmed to 65°C and its concentration was quantified using a Qubit 3.0 fluorometer. Illumina Tn-seq library construction was as described^28^. Briefly, gDNA was sonicated and ligated with common adapters, and for each fragment deriving from a Tn insertion in the genome, a sequence containing a portion of the transposon and a portion of its genomic context (the Tn-genome junction) was amplified using one primer homologous to a region in the transposon, and another primer homologous to a region in the adapter. The indexed adapter-specific primer was CAAGCAGAAGACGGCATACGAGATNNNNNNGTGACTGGAGTTCAGACGTGTGCTCTTCCG ATCT, where the six N’s are a unique index used for multiplexing multiple libraries onto the same Hiseq sequencing lane, and the transposon specific primer was ATGATACGGCGACCACCGAGATCTACACTCTTTCCCTACACGACGCTCTTCCGATCT NNNNNNAGCAATATTTCAAGAATGCATGCGTCAAT, where N’s are random nucleotides. Amplification used Jumpstart polymerase (Sigma) and the following cycling protocol: 94°C-2 min, [94°C-30 sec, 65°C-20 sec, 72°C-30 sec] X 25, 72°C-10 min. Sequencing of single-end reads of 150 bp was done over eight lanes on a HiSeq 2500 at the Joint Genome Institute (Walnut Creek, CA). Reads sequenced per library are reported in Table S2.

### Tn-seq read-mapping and data analysis

For analysis of data from the sequencing of Tn insertion sites in pools of hemizygotes, we first searched each read for a string corresponding to the last 20 base pairs of the left arm of the piggyBac transposon sequence, allowing up to two mismatches. For each Tn-containing read, we then identified the genomic location of the sequence immediately downstream of the Tn insertion site, which we call the genomic context of the insertion, by mapping with BLAT (minimum sequence identity = 95, tile size = 12) against a hybrid reference genome made by concatenating amended *S. cerevisiae* DBVPG1373 and *S. paradoxus* Z1 genomes (see below). These genomic-context sequence fragments were of variable length; any case in which the sequence was shorter than 50 base pairs was eliminated from further analysis, as was any case in which a genomic-context sequence mapped to more than one location in the hybrid reference. The resulting data set thus comprised reads containing genomic-context sequences specifically mapping to a single location in either *S. cerevisiae* DBVPG1373 or *S. paradoxus* Z1, which we call usable reads. For a given library, given a cohort of usable reads whose genomic-context sequence mapped to the same genomic location, we inferred that these reads originated from clones of a single mutant with the Tn inserted at the respective site, which we call an insertion. In cases where the genomic-context sequences from reads in a given library mapped to positions within 3 bases of each other, we inferred that these all originated from the same mutant genotype and combined them, assigning to them the position corresponding to the single location to which the most reads mapped among those combined. For a given insertion thus defined, we considered the number of associated reads n_insert_ as a measure proportional to the abundance of the insertion clone in the cell pellet whose gDNA was sequenced. To enable comparison of these abundances across samples, we tabulated the total number of usable reads n_pellet_ from each cell pellet, took the average of this quantity across pellets, <n_pellet_>, and multiplied each n_insert_ by <n_pellet_>/n_pellet_ to yield a_insert_, the final normalized estimate of the abundance of the insertion clone in the respective pellet. For any insertions that were not detected in a given pellet’s library (n_insert_ = 0) but detectable in another library of the data set, we assigned n_insert_ = 1.

We evaluated, from the mapped genomic-context sequence of each insertion, whether it fell into a gene according to the *S. cerevisiae* and *S. paradoxus* genome annotations^17,29^, and we retained for further analysis only those insertions that fell into genes that were syntenic in the two species. For each such insertion, for each biological replicate corresponding to a selection culture (at 28°C or 39°C), we averaged the normalized abundances a_insert_ across technical replicates, yielding a single abundance estimate <a_insert_>_technical_ for the biological replicate. We then calculated the mean of the latter quantities across all biological replicates of the selection, to yield a final abundance estimate for the insertion in this selection, <a_insert_>_total_. Likewise, for each insertion and selection experiment we calculated CV_insert,total_, the coefficient of variation of <a_insert_>_technical_ values across biological replicates.

To use Tn-seq data in reciprocal hemizygosity tests, we considered for analysis only genes annotated with the same (orthologous) gene name in the *S. cerevisiae* and *S. paradoxus* reference genomes. For each insertion, we divided the <a_insert_>_total_ value from the 39°C selection by the analogous quantity from the 28°C selection and took the log_2_ of this ratio, which we consider to reflect thermotolerance as measured by RH-seq. For each gene in turn, we used a two-tailed Mann-Whitney U test to compare thermotolerance measured by RH-seq between the set of insertions falling into the *S. cerevisiae* alleles of the gene, against the analogous quantity from the set of insertions falling into the *S. paradoxus* allele of the gene, and we corrected for multiple testing using the Benjamini-Hochberg method.

We tabulated the number of inserts and genes used as input into the reciprocal hemizygote test, and the number of top-scoring genes emerging from these tests, under each of a range of possible thresholds for coverage and measurement noise parameter values (Figure S5). We used in the final analysis the parameter-value set yielding the most extensive coverage and the most high-significance hits: this corresponded to insertions whose abundances had, in the data from at least one of the two selections (at 28°C or 39°C), CV_insert,total_ ≤ 1.5 and <a_insert_>_total_ ≥ 1.1, and genes for which this high-confidence insertion data set contained at least 5 insertions in each species’ allele. This final data set comprised 110,678 high-quality insertions (Table S3) in 3416 genes (Table S4).

### Amended reference genome construction

We generated reference genomes for *S. cerevisiae* strain DBVPG1373 and *S. paradoxus* strain Z1 as follows. Raw genome sequencing reads for each strain were downloaded from the SGRP2 database (ftp://ftp.sanger.ac.uk/pub/users/dmc/yeast/SGRP2/input/strains). Reads were aligned using bowtie2^30^ with default options; DBVPG1373 reads were aligned to version R64.2.1 of the reference sequence of the *S. cerevisiae* type strain S288C (Genbank Assembly Accession GCA_000146045.2), and Z1 reads were aligned to the *S. paradoxus* strain CBS432 reference sequence^31^. Single nucleotide variants (SNPs) were called using a pipeline of samtools^32^, bcftools and bgzip, and were filtered for a quality score (QUAL) of >20 and a combined depth (DP) of >5 and either <65 (*S. cerevisiae*) or <255 (*S. paradoxus*). We then amended each reference genome with the respective filtered SNPs: we replaced the S288C allele with that of DBVPG1373 at each filtered SNP using bcftools’ consensus command with default options (42,983 base pairs total), and amendment of the CBS432 sequence was carried out analogously using Z1 alleles (15,126 base pairs total).

### Targeted-deletion hemizygote construction by homologous recombination

A given targeted hemizygote for each RH-seq hit gene except *TAF2* was generated in the *S. cerevisiae* DBVPG1373 x *S. paradoxus* Z1 hybrid (JR507) by knocking out the allele of the gene from one species via homologous recombination with KANMX as described^33^ with 70 base pairs of homology on the 5’ and 3’ ends of the cassette; checking was via diagnostic PCR. Two or more independent transformants were isolated and phenotyped for each hemizygote genotype (Table S1).

### Construction of allele replacement and targeted hemizgyote strains with Cas9

At each RH-seq hit gene, we constructed strains in wild-type homozygous diploid *S. cerevisiae* DBVPG1373 in which both copies of the endogenous allele were replaced by the allele from an *S. paradoxus* isolate (Z1 for Figures 3, S8, and S9, and other strains as indicated for Figure S10), and likewise for replacement of alleles from *S. cerevisiae* (DBVPG1373 in Figures 3, S8, and S9, and other strains as indicated for Figure S10) into *S. paradoxus* Z1. We call each such strain an allele-replacement strain, and each was constructed using a dual-guide Cas9 transgenesis method^34^ in which a linear PCR fragment from the donor species is incorporated into the recipient genome by homology-directed repair of two chromosomal double-strand breaks induced by Cas9. Briefly, for each allele of each gene, we designed two guide RNAs for double-strand breaks by Cas9: one guide targeted a position ~1000 base pairs 5’ to the coding start or at the 3’ end of the closest upstream gene, whichever was closer, and the other guide targeted the region of the coding stop. The precise cut site of each was chosen to contain an NGG immediately downstream of variants between the *S. cerevisiae* and *S. paradoxus* strains, to avoid re-cutting of the donor allele by Cas9 after it had been introduced into the recipient strain. We cloned the two guide RNAs, a KANMX cassette, and the gene encoding the *S. pyogenes* Cas9 protein into a single plasmid as described^34^. The resulting plasmid was propagated in DH5a *E. coli* and miniprepped with a column miniprep kit (Qiagen). Separately, to generate the fragment to be used as the donor for DNA repair after Cas9 cutting, we PCR-amplified the respective region from the donor strain, with primers whose 5’-most 70 base pairs were homologous to the recipient and whose 3’-most 20 base pairs were homologous to the donor, except in the case of transgenesis using *S. cerevisiae* donors other than DBVPG1373 or *S. paradoxus* donors other than Z1 (for Figure S10), for which the 3’-most 20 base pairs were homologous to DBVPG1373 or Z1, respectively. The homology region at the 5’ end of the gene ended at most 31 base pairs upstream of the 5’ cut site, and the homology region at the 3’ end of the gene started at most 33 base pairs downstream of the 3’ cut site. The donor fragment product was purified with a column kit (Qiagen) and ethanol-precipitated. We then simultaneously transformed, using the lithium acetate method, the donor fragment and dual-guide Cas9 plasmid into the recipient strain, using donor:acceptor ratios of 0.38:10 to 1:5, with 0.5-26 ug of plasmid. In this transformation, heat shock was for 20-30 minutes at 39-42°C in transformations of *S. cerevisiae* DBVPG1373, and 10-20 minutes at 37-39°C for transformations of *S. paradoxus* Z1. Transformants were plated on YPD+G418 (300 μg/mL) to select for cells that retained the plasmid. From this selection we patched single colonies onto YPD without G418, under the expectation that by the time a lawn came up for each patch, its cells would have lost the Cas9 plasmid. Each such strain was Sanger-sequenced at the junctions of the recipient and donor sequence. Positive patches were streaked to single colonies on YPD plates, and cells from each such colony were used to inoculate a patch on a YPD plate and, separately, to inoculate a patch on a YPD+G418 plate. Those colonies whose patches grew on the former but not on the latter were inferred to be cured of the plasmid and stored at −80°C. For all genes except *DYN1*, 2-3 such strains from each transformation were retained for thermotolerance assays and underwent Sanger sequencing of the entire locus to determine the exact swapped region (Table S1). For *DYN1* allele-replacement in *S. paradoxus* Z1, the Cas9-based strategy yielded a single verified clone in which the *S. paradoxus* Z1 allele of *DYN1* was replaced by that of *S. cerevisiae* DBVPG1373, and likewise for *DYN1* allele-replacement in *S. cerevisiae* DBVPG1373. In each case, we mated the single swap clone to a wild-type of the respective species background, confirmed heterozygosity of the resulting diploid via allele-specific diagnostic PCR at the *DYN1* locus, sporulated, and dissected tetrads, allowing each spore to autodiploidize and grow up as a homozygote; we retained from one such tetrad the two spores that were homozygous at the *DYN1* locus for the swapped allele, as confirmed by sequencing, and stored these at −80°C.

Targeted-deletion hemizygote strains for *TAF2* were generated by knocking out the *S. cerevisiae* or the *S. paradoxus* allele in the interspecific hybrid (JR507) using the above methods for Cas9 cutting and repair, with the following differences. To generate the fragment to be used as the donor for DNA repair after Cas9 cutting, we PCR-amplified the NATMX cassette from pBC713 (a gift from John Dueber, constructed as in^35^) using primers whose 5’-most 70 base pairs were homologous to the recipient and whose 3’-most 20 base pairs were homologous to the cassette. The homology region at the 5’ end of the gene ended 22 base pairs outside the 5’ cut site (which was upstream of coding start), and the homology region at the 3’ end of the gene started 33 base pairs outside the 3’ cut site. Positive strains were confirmed by PCR. Two independent transformants were isolated and phenotyped for each genotype (Table S1).

### Growth assays

#### Growth measurements of wild-type SGRP strains

For the growth timecourse of a given wild-type, purebred homozygote isolate from the SGRP collection at 28°C, it was streaked from a −80°C freezer stock onto a YPD agar plate and incubated at 26°C for 3 days. A single colony was inoculated into liquid YPD and grown for 24 hours at 28°C with shaking at 200 rpm. This culture was back-diluted into YPD at an OD_600_ of ~0.05 and grown for an additional 5.5 hours at 28°C, 200 rpm, until reaching logarithmic phase. We transferred cells from each such pre-culture, and YPD, to each of 11 replicate wells of a 96-well plate, with volumes sufficient to yield a total volume of 150 μL per well at an OD_600_ of 0.02. The plate was covered with a gas-permeable membrane (Sigma) and incubated with orbital shaking in an M200 plate reader (Tecan, Inc.) at 28°C for 24 hours. For curves in Figure S1A, measurements for optical density at 595nm (OD_595_) were taken every 30 minutes and for each timepoint, the average was taken across replicate wells. To subtract background OD_595_ for the resulting curve, we tabulated the mean of the five lowest values from all datapoints, excluding the first two, and subtracted this value from that of each timepoint, setting any negative value to 0. To smooth the resulting curve, we first replaced each timepoint measurement by its average with those of the timepoints immediately before and after it; then, for any timepoint whose measurement was not greater than or equal to the previous one, we set it to be equal to that previous data point. For Figure S1B, the efficiency for a given growth curve (from a single well) was calculated as the difference between the OD_595_ measured at the last four smoothed and averaged data points and that of the first four smoothed and averaged data points. Efficiencies from all of the wells from every *S. cerevisiae* isolate were combined and compared to efficiencies from all of the wells for every *S. paradoxus* isolate in a two-sample two-tailed *t*-test.

For the growth timecourse of a given SGRP strain at 39°C (Figure 1A), we used a large-volume flask growth paradigm to avoid the influence of plate effects on growth measurements at high temperature in the incubated microplate reader, as follows. Each strain was streaked from a −80°C freezer stock onto a YPD agar plate and incubated at 26°C for 3 days. A single colony of a given strain was inoculated into liquid YPD and grown for 24 hours at 28°C with shaking at 200 rpm. Each of these cultures was back-diluted into YPD at an OD_600_ of 0.05 and grown for an additional 5.5-7.5 hours at 28°C, shaking at 200 rpm, until reaching logarithmic phase. We transferred cells from each such pre-culture, and YPD, to a glass 250 mL flask at the volumes required to attain an OD_600_ of 0.05 in 100 mL YPD, and incubated it at 39°C with shaking at 200 rpm. OD_600_ readings were taken every ~2 hours for ~18 hours. Figure 1A reports representative data from one of three such independent timecourse experiments. For curve fits, we used the getInitial and SSlogis functions in R to estimate starting values for the parameters of the logistic equation, and the nls function to fit the final parameters.

For efficiency measurements of a given SGRP isolate at 39°C in the large-volume format (Figure 1B), it was streaked from a −80°C freezer stock onto a YPD agar plate and incubated at 26°C for 3 days. Two single colonies of each isolate were each inoculated into liquid YPD and grown for 24 hours at 28°C with shaking at 200 rpm to generate two replicate pre-cultures. Each was back-diluted into YPD at an OD_600_ at 600 nm of 0.05 and grown for an additional 5.5 hours at 28°C, shaking at 200 rpm, until reaching logarithmic phase. The two pre-cultures were each again back-diluted into YPD in 1-inch diameter glass tubes with a target OD_600_ of 0.05; the actual OD_600_ of each was measured, after which it was grown at 39°C with shaking at 200rpm for 24 hours, and OD_600_ was measured again. The efficiency for each replicate was calculated as the difference between these final and initial OD_600_ values. The pipeline from inoculation off solid plates through pre-culture, two back-dilutions, and growth at 39°C we refer to as a day’s growth experiment. For each day’s experiments, we calculated the average efficiency across the replicates of each isolate <e_strain_>. We carried out two days’ worth of replicate growth experiments for each isolate. For a given species, we used the complete cohort of measurements of <e_strain_> from all isolates of each species, across all days, as input into a two-sample, two-tailed t-test to evaluate whether the suite of e_strain_ values across isolates of *S. cerevisiae* was significantly different from the analogous set of values from *S. paradoxus*.

#### Growth measurements of targeted-deletion hemizygotes and allele-replacement strains at 28°C

For efficiency measurements of a given targeted-deletion hemizygote or allele-replacement strain at 28°C (Figures S7, S8 and S10), pre-culture and plate reader assays were as for wild-type SGRP strains, except that 6 or more replicate wells were cultured per strain. 2-3 independently isolated targeted-deletion hemizygotes or allele-replacement strains (Table S1) were assayed for each genotype. Each timecourse of targeted-deletion hemizygote or allele-replacement strains also included the wild-type hybrid (JR507) or parent (*S. cerevisiae* DBVPG1373 or *S. paradoxus* Z1), respectively, with pre-culture and replication as above. Efficiency for a given growth curve (from a single well) was calculated as the difference between the OD_600_ measured at the last four smoothed and averaged datapoints and that of the first four smoothed and averaged datapoints, with smoothing and averaging as detailed above. For Figure S7, relative efficiency for a given well of a given targeted-deletion hemizygote strain at 28°C was tabulated as its efficiency divided by that of the average of all replicate wells of the wild-type hybrid (JR507) grown in the same experiment. For a given gene, we used the complete cohort of these measurements, from all isogenic hemizygotes, as input into a two-sample, two-tailed t-test to evaluate whether the relative efficiency of the strain in which the *S. cerevisiae* allele was knocked out was lower than the analogous quantity from the strain in which the *S. paradoxus* allele was knocked out. In Figures S8 and S10, allele-replacement strains for a given gene were analyzed analogously, with the relative efficiency calculated against the respective wild-type parent, and with a one-sample, two-tailed t-test to evaluate whether the relative efficiency was significantly different from 1.

#### Growth measurements of targeted-deletion hemizygotes and allele-replacement strains at 39°C

For efficiency measurements of a given targeted-deletion hemizygote strain at 39°C in the large-volume format (Figure 2B), each strain was streaked from a −80°C freezer stock onto a YPD agar plate and incubated at 26°C for 3 days. Two single colonies of a given strain were each inoculated into liquid YPD and grown for 24 hours at 28°C with shaking at 200 rpm. Each such pre-culture at stationary phase, or a log-phase outgrowth of it (which we used in the case of *DYN1* and *TAF2:* the pre-culture and YPD were added at the volumes required to attain an OD_600_ of 0.05 and grown for an additional 5.5 hours at 28°C, shaking at 200 rpm, until the culture reached logarithmic phase) was used to inoculate YPD in 1-inch diameter glass culture tubes with a target cell density corresponding to an OD_600_ of 0.05. The actual OD_600_ of each was measured, after which it was grown at 39°C with shaking at 200 rpm for 24 hours, and OD_600_ was measured again. The efficiency for each such replicate was then calculated as the difference between the final and initial OD_600_ values. The pipeline from inoculation off solid plates through pre-culture, back-dilution, and growth at 39°C we refer to as a day’s growth experiment for a targeted-deletion homozygote. In each such experiment, 2-3 independently isolated targeted-deletion hemizygotes of a given gene in each direction were all assayed on the same day, alongside the wild-type hybrid parent (JR507) with replicate structure and methods as above. From each day’s experiments, we calculated the average efficiency across the replicates of the wild-type hybrid <e_hybrid_>, and we used this quantity to normalize the efficiency e_hemizyg_ measured for each replicate of each hemizygote strain assayed on that day. Thus, the final observable used for analysis for each replicate on a given day was e_hemizyg_/<e_hybrid_>. We carried out 2-3 days’ replicate growth experiments for each gene’s hemizygotes. For a given gene, we used the complete cohort of these measurements of e_hemizyg_/<e_hybrid_>, from all days and all isogenic hemizygotes, as input into a two-sample, one-tailed t-test to evaluate whether e_hemizyg_/<e_hybrid_> of the strain in which the *S. cerevisiae* allele was knocked out was lower than the analogous quantity from the strain in which the *S. paradoxus* allele was knocked out.

For growth measurements of a given allele-replacement strain at 39°C in the large-volume format (Figure 3, S7, S8 and S10), each strain was streaked from a −80°C freezer stock onto a YPD agar plate and incubated at 26°C for 3 days. 1-2 single colonies of each strain were each inoculated into liquid YPD and grown for 24 hours at 28°C with shaking at 200 rpm to generate 1-2 replicate pre-cultures. Each was back-diluted into YPD at an OD_600_ of 0.05 and grown for an additional 5.5 hours at 28°C, shaking at 200 rpm, until reaching logarithmic phase. Each pre-culture were each again back-diluted into YPD in 1-inch diameter glass tubes with a target OD_600_ of 0.05 (for experiments using a single pre-culture, it was now split into two replicate precultures, each of the same OD_600_); the actual OD_600_ of each was measured, after which it was grown at 39°C with shaking at 200rpm for 24 hours, and OD_600_ was measured again. The efficiency for each replicate was calculated as the difference between these final and initial OD_600_ values. The pipeline from inoculation off solid plates through pre-culture, two back-dilutions, and growth at 39°C we refer to as a day’s growth experiment for an allele-swap strain. In each such experiment, 2-3 independently isolated allele-swap strains targeting a given gene in a given background were all assayed on the same day, alongside the respective wild-type background strain (*S. cerevisiae* DBVPG1373 or *S. paradoxus* Z1) with replicate structure and methods as above. For each day’s experiments, we calculated the average efficiency across the replicates of the wild-type parent <e_parent_>, and we used this quantity to normalize the efficiency e_swap_ measured for each replicate assayed on that day of each allele-swap strain in the respective background. Thus, the final measurement used for analysis for each replicate on a given day was e_swap_/<e_parent_>. We carried out 2-3 days’ worth of replicate growth experiments for each gene’s allele-swap strains. For a given gene in a given background, we used the complete cohort of measurements of e_swap_/<e_parent_> from all days and all allele-swap strains as input into a one-sample, one-tailed t-test to evaluate whether e_swap_/<e_parent_> was significantly different from 1. For swaps of the *S. cerevisiae* allele of a given gene into *S. paradoxus*, we tested whether e_swap_/<e_parent_> was greater than 1 (*i.e*. that the swap strain grew better at 39°C than did its parent), and for swaps of the *S. paradoxus* allele of a given gene into *S. cerevisiae*, we tested whether e_swap_/<e_parent_> was less than 1 (*i.e*. that the swap strain grew worse at 39°C than its parent).

#### Testing survival of wild-type parent strains at 39°C

To test the survival of heat-treated *S. paradoxus* Z1 and *S. cerevisiae* DBVPG1373 (Figure S2), each strain was streaked from −80°C freezer stocks onto YPD agar plates and incubated at 26°C for 3 days. 1-2 single colonies of each parent were inoculated into liquid YPD and grown for 24 hours at 28°C with shaking at 200 rpm to create 1-2 replicate pre-cultures. After 24 hours, we transferred cells from each pre-culture, and YPD, to each of 2-4 tubes at the volumes required to attain an OD_600_ of 0.05 in 11 mL YPD. Two tubes were incubated at 28°C and two at 39°C, all with shaking at 200 rpm for 24 hours. OD_600_ of each was measured; for each culture grown at 28°C, 100 μL of a 1.0×10^−5^ serial dilution was plated to YPD, and for cultures grown at 39°C, *S. paradoxus* and *S. cerevisiae* were serially diluted to 10^−1^ and 5.0×10^−5^, respectively. Plates were incubated at 26°C for three days until single colonies appeared. Colonies on each plate were counted, from which we tabulated the colony forming units per mL of culture plated, and normalized by the optical density of the original plated culture. The pipeline from pre-culture through treatment and colony counting we refer to as one day’s worth of experiments. We used the results of two days’ worth of experiments (a total of four for each species and temperature) as input into an ANOVA with species and temperature as factors, and took the p-value for the interaction between the factors as the estimate of the significance of the difference between species in the effect of temperature (of the 24-hour liquid culture) on cells’ ability to grow into colonies (at the permissive temperature).

### Microscopy and quantification

For microscopy of *S. cerevisiae* DBVPG1373 and *S. paradoxus* Z1 (Figure S3), each species was streaked from a −80°C freezer stock onto a YPD agar plate and incubated at 26°C for 3 days. Two single colonies of each strain were each inoculated into liquid YPD and grown for 24 hours at 28°C with shaking at 200 rpm to generate two replicate pre-cultures. Each was then back-diluted into YPD at an OD_600_ of 0.05; one was grown at 39°C with shaking at 200rpm for 24 hours, and the other was grown at 28°C with shaking at 200rpm for 24 hours. After the 24 hour growth period, 0.5 OD units of each culture were harvested through centrifugation and incubated in 66.5% ethanol for 1-4 hours at room temperature. Each sample was washed twice with 1X Dulbecco’s Phosphate Buffered Saline (DPBS, Gibco), resuspended in 0.5 mL of 1X DPBS, and vortexed for 15 seconds on high. 5 μL of each sample was transferred to an agarose pad made with 1% agarose and YPD. We observed samples on a Zeiss Axio Observer inverted bright-field microscope at 100X magnification. Images were taken using a Hamamatsu ORCA-Flash4.0 digital camera and visualized using ZEN software for image analysis. The exposure of each image was set automatically through ZEN, and brightness was adjusted using the “Min/Max” adjustment for black and white light. The pipeline from inoculation off solid plates through pre-culture, growth at 39°C or 28°C, fixation, and imaging we refer to as a day’s experiment. We carried out 2 days’ worth of experiments for each species, yielding a total of 17-21 images. In each image, free-floating cells were scored manually as singlets or those with a small, medium, or large bud (for a total of 31-151 scored cells per species and temperature).

The proportion of scored cells with large buds was tabulated for each day’s experiment for each species and temperature. We used these proportions (a total of two for each species and temperature) as input into an ANOVA with species and temperature as factors, and took the p-value for the interaction between the factors as the estimate of the significance of the difference between species in the effect of temperature on the proportion of cells with large buds.

### Sequence analysis

#### D_xy_ *analysis*

To assess the divergence between *S. cerevisiae* and *S. paradoxus* at loci mapped by RH-seq, we used the *D_xy_* statistic^36^, the average number of pairwise differences between all *S. cerevisiae* strains and *S. paradoxus*, normalized for gene length, as follows. We downloaded *S. cerevisiae* genomic sequences from the following sources: YJM978, UWOPS83-787, Y55, UWOPS05-217.3, 273614N, YS9, BC187, YPS128, DBVPG6765, YJM975, L1374, DBVPG1106, K11, SK1, 378604X, YJM981, UWOPS87-2421, DBVPG1373, NCYC3601, YPS606, Y12, UWOPS05-227.2, and YS2 from http://www.yeastrc.org/g2p/home.do; Sigma1278b, ZTW1, T7, and YJM789 from http://www.yeastgenome.org/; and RM11 from NCBI (accession PRJNA13674). For each strain, we extracted the coding sequence of each gene in turn, and we downloaded the *S. paradoxus* reference sequence for each orthologous coding region from^17^. Sequences were aligned using MAFFT^37^ with default settings. Alignments that did not contain a start and stop codon, or those that contained gaps at greater than 40% of sites were considered poor quality and discarded. We tabulated *D_xy_* for each gene. To evaluate whether the eight RH-seq hit genes were enriched for elevated *D_xy_*, we first tabulated <D_xy_>_true_, the mean value across the eight RH-seq hit genes. We then sampled eight random genes from the set of 3416 genes tested by RH-seq; to account for biases associated with lower rates of divergence among essential genes, the resampled set contained six essential genes and two non-essential genes, mirroring the breakdown of essentiality among the RH-seq hits. Across this random sample we tabulated the mean *D_xy_*, <D_xy_>_resample_. We repeated the resampling 5000 times and used as an empirical p-value the proportion of resamples at which <D_xy_>_resample_ ≤ <D_xy_>_true_.

#### Phylogenetic analysis

We downloaded orthologous protein coding regions for the type strains of *S. cerevisiae, S. paradoxus*, and an outgroup, *S. mikatae*, from^17^. For each gene for which ortholog sequences were available in all three species, we aligned the sequences with PRANK^38^ utilizing the “-codon” option for codon alignment. These alignments were used as input into the codeml module of PAML^39^, which was run assuming no molecular clock and allowing omega values to vary for each branch in the phylogeny. From the resulting inferences, we tabulated the branch length on the *S. cerevisiae* lineage for each gene. To evaluate whether sequence divergence of the eight RH-seq hit genes showed signatures of rapid evolution along the *S. cerevisiae* lineage, we used the resampling test detailed above.

### Locus effect size

Locus effect sizes in Figure S9 were computed from the data in Figures 2B and 3 of the main text as follows. For analyses in the hybrid background, for a given locus we calculated *m_S.par_*, the mean of all replicate measurements of e_hemizyg_/<e_hybrid_> of hemizygote strains lacking the *S. paradoxus* allele. We took, as independent estimates of the locus effect size (each corresponding to an open circle on the respective grey bar in Figure S9), each measurement of e_hemizyg_/<e_hybrid_> of a hemizygote strain lacking the *S. cerevisiae* allele, as a ratio against *m_S.par_*. For analyses in the *S. cerevisiae* DBVPG1373 background, for a given locus we used each measurement of e_swap_/<e_parent_> as an independent estimate of the locus effect size (each corresponding to an open circle on the respective orange bar in Figure S9). For analyses in the *S. paradoxus* Z1 background, for a given locus we used each measurement of <e_parent_>/e_swap_ as an independent estimate of the locus effect size (each corresponding to an open circle on the respective blue bar in Figure S9).

### Data availability

RH-seq data will be deposited in the Sequence Read Archive (SRA) upon publication.

### Code availability

Custom Python and R scripts used for RH-seq data analysis are available upon request.

## End Notes

**Supplementary Information** is available in the online version of the paper.

## Acknowledgements

The authors thank Faisal AlZaben, Anna Flury, Gina Geiselman, Justin Hong, Jake Kim, Matt Maurer, and Luke Oltrogge for technical assistance, Dave Savage for his generosity with microscopy resources, and Ben Blackman, Sam Coradetti, Avi Flamholz, Vinny Guacci, Doug Koshland, Chris Nelson, and Arjun Sasikumar for discussions. This work was supported by R01 GM120430-A1 and by Community Sequencing Project 1460 to RBB at the U.S. Department of Energy Joint Genome Institute, a DOE Office of Science User Facility. The work conducted by the latter was supported by the Office of Science of the U.S. Department of Energy under Contract No. DE-AC02-05CH11231.

## Author Contributions

R.B.B. and J.I.R. developed the project design; C.V.W., J.I.R., R.H. and J.C. performed experiments; C.V.W. and J.I.R. and analyzed the data; J.M.S. contributed to the development of mutagenesis and sequencing methods; A.P.A. contributed mutagenesis and sequencing reagents; I.V.G. provided technical assistance with sequencing; R.B.B. and C.V.W wrote the manuscript with input from all authors.

## Author Information

The authors do not declare any competing financial interests. Correspondence and request for materials should be addressed to R.B.B. and J.I.R.

